# H3K4me3 is neither instructive for, nor informed by, transcription

**DOI:** 10.1101/709014

**Authors:** Struan C Murray, Philipp Lorenz, Françoise S Howe, Meredith Wouters, Thomas Brown, Shidong Xi, Harry Fischl, Walaa Khushaim, Joseph Regish Rayappu, Andrew Angel, Jane Mellor

## Abstract

H3K4me3 is a near-universal histone modification found predominantly at the 5’ region of genes, with a well-documented association with gene activity. H3K4me3 has been ascribed roles as both an instructor of gene expression and also a downstream consequence of expression, yet neither has been convincingly proven on a genome-wide scale. Here we test these relationships using a combination of bioinformatics, modelling and experimental data from budding yeast in which the levels of H3K4me3 have been massively ablated. We find that loss of H3K4me3 has no effect on the levels of nascent transcription or transcript in the population. Moreover, we observe no change in the rates of transcription initiation, elongation, mRNA export or turnover, or in protein levels, or cell-to-cell variation of mRNA. Loss of H3K4me3 also has no effect on the large changes in gene expression patterns that follow galactose induction. Conversely, loss of RNA polymerase from the nucleus has no effect on the pattern of H3K4me3 deposition and little effect on its levels, despite much larger changes to other chromatin features. Furthermore, large genome-wide changes in transcription, both in response to environmental stress and during metabolic cycling, are not accompanied by corresponding changes in H3K4me3. Thus, despite the correlation between H3K4me3 and gene activity, neither appear to be necessary to maintain levels of the other, nor to influence their changes in response to environmental stimuli. When we compare gene classes with very different levels of H3K4me3 but highly similar transcription levels we find that H3K4me3-marked genes are those whose expression is unresponsive to environmental changes, and that their histones are less acetylated and dynamically turned-over. Constitutive genes are generally well-expressed, which may alone explain the correlation between H3K4me3 and gene expression, while the biological role of H3K4me3 may have more to do with this distinction in gene class.

## Introduction

Histones are universally recognised as an integral component of eukaryotic gene regulation. Histone octamers bind DNA to form nucleosomes, winding the DNA and changing its binding surface, fundamentally altering its accessibility and structure. The extruding tails of the histone proteins can also undergo extensive covalent modification. These histone modifications can alter nucleosome structure and recruit proteins involved in gene regulation, and so can play a role in the modulation of gene expression. However, the extent to which these modifications are intrinsic or essential to physiological gene expression and regulation is not well understood.

Arguably the most well-studied histone modification is the trimethylation of lysine 4 of histone H3 (H3K4me3). Inputting the search term ‘H3K4me3’ into Pubmed yields nearly 1800 hits, while ‘H3K27’ yields 1315 hits, ‘H3K36me3’ 380, with other histone residues and modifications yielding far less. One likely reason for H3K4me3’s popularity is its well-established association with gene expression (Santos-Rosa et al., 2002). The levels of H3K4me3 at an averaged or ‘meta’ gene show a pronounced peak immediately downstream of the transcription start site (TSS) that rapidly tails off (Pokholok et al., 2005). The height of this peak is well correlated with the level of gene expression (Howe et al., 2017).

As a result of this association, H3K4me3 is often referred to as an ‘activating’ histone modification, with a rise in H3K4me3 expected to cause a subsequent rise in gene expression. However, whether the association of H3K4me3 with gene expression is the result of a *causal* relationship whereby H3K4me3 instructs transcription has never been firmly established (Howe et al., 2017).

Like numerous other histone modifications, H3K4me3 is capable of recruiting proteins to chromatin, which could result in the upregulation of transcription. Notably, H3K4me3 has been shown to interact with the general transcription factor TFIID, meaning it could serve to stabilise the formation of the pre-initiation complex (Lauberth et al., 2013; Vermeulen et al., 2007). H3K4me3 binds to a number of different characteristic protein folds, including the PHD finger, the Tudor domain, and the chromodomain (Ruthenburg et al., 2007). Such folds can be found in a range of chromatin remodelling and histone modifying factors, and so could reasonably be expected to mediate a change in transcription in response to changing levels of H3K4me3.

There is also evidence that H3K4me3 can itself result as a consequence of transcription. For example, Vastenhouw et al. (2010) reported that the bulk of H3K4me3 occurs after the start of transcription during zygotic activation in zebra fish. Trimethylation of H3K4me3 has been shown to depend upon the phosphorylation of the C-terminal domain of RNA Polymerase II (RNAPII), specifically at Serine 5 (Lee and Skalnik, 2008; Ng et al., 2003). This phosphorylation is deposited by TFIIH, with a sharp peak present at the 5’ end of the gene body (Ng et al., 2003). This could in turn explain why H3K4me3 is predominantly found at the 5’ end of genes. Indeed, covalently fusing the Set1 protein required for H3K4 methylation to RNAPII results in an extended H3K4me3 peak into the body of highly expressed genes (Soares et al., 2017). What might the purpose of this H3K4me3 deposition be? Ng et al. (2003) proposed that H3K4me3 might act as a sort of transcriptional memory, based on the observation that H3K4me3 levels at the *GAL10* gene fall away much more gradually than RNAPII following gene inactivation – the implication being that the hypothesised activatory nature of H3K4me3 might allow for a recently repressed gene to become quickly active again.

H3K4 methylation is performed by highly conserved enzymes containing the SET methyltransferase domain. These enzymes are present within complexes with other regulatory proteins - the SET1 complexes in *S. cerevisiae* and humans. One of these subunits – Spp1 and CFP1 in *S. cerevisiae* and humans respectively – is necessary for maximal levels of H3K4me3; ablation of these proteins specifically and drastically reduce H3K4me3 with little effect on H3K4me1 and H3K4me2 levels (Morillon et al., 2005; Schneider et al., 2005; Thomson et al., 2010). In *S. cerevisiae*, deletion of Spp1 reduces global levels of H3K4me3 to ~25% (Howe et al., 2017; Sommermeyer et al., 2013).

We sought to test the hypothesis that H3K4me3 is universally instructive for gene expression. To achieve this, we took advantage of the above observation that deletion of Spp1 reduces H3K4me3 without affecting related marks in *S. cerevisiae*. We assessed the effects of lowered H3K4me3 at multiple levels of gene expression by utilising a range of experimental and modelling-based approaches. By determining the rates of transcription initiation, elongation and mRNA degradation, the levels of steady-state mRNA and protein, and other gene parameters like expression noise, we find that H3K4me3 is *not* instructive for gene expression on a genome-wide scale. Conversely, by depleting RNA Polymerase II from the nucleus, we find that the pattern and total level of H3K4me3 genome-wide remains unchanged after an hour, while changes in H3K4me3 associated with environmental stress are small and unrelated to larger changes in transcription. Taken together, our findings demonstrate that transcription and H3K4me3, though well correlated, do not inform and are not required to maintain the levels of the other.

## Results

### H3K4me3 predominantly marks highly expressed, environmentally insensitive genes

We wanted to ask whether H3K4me3 modulates gene expression at any level. To this end we deleted the *SPP1* gene, in order to abrogate H3K4me3 levels genome-wide. We used ChIP-seq to assess H3K4me3 levels in both wild-type and *spp1Δ* strains. Importantly, we spiked-in our *S. cerevisiae* cells with known quantities of *S. pombe* prior to sequencing, so that our data could be properly normalised, and so allowing for direct comparisons between strains.

Our H3K4me3 ChIP-seq data was in good agreement with existing genome-wide datasets, with the same distinctive 5’ peak typical of H3K4me3 meta-gene profiles (Fig 1A) (Pokholok et al., 2005). As expected, levels of H3K4me3 are drastically reduced in *spp1Δ* (Fig 1A-B). The total genome-wide read-count, after normalising to account for the *S. pombe* spike-in, was ~25% of what it was in wild-type (Fig 1B). This is a similar reduction to what we observe globally by western blot (Fig 1C) and at the *FMP27* gene by RT-PCR (Fig D). This reduction is fairly uniform genome-wide, evidenced by the fact that despite this huge reduction, the levels of H3K4me3 in *spp1Δ* still correlate well genome-wide with the levels in wild-type (Fig 1E-G, spearman correlation coefficient = 0.68). This suggests that the loss of Spp1 results in a substantial reduction in Set1’s ability to produce H3K4me3, without completely abolishing its catalytic activity or substantially affecting its targeting.

**Figure 1.**
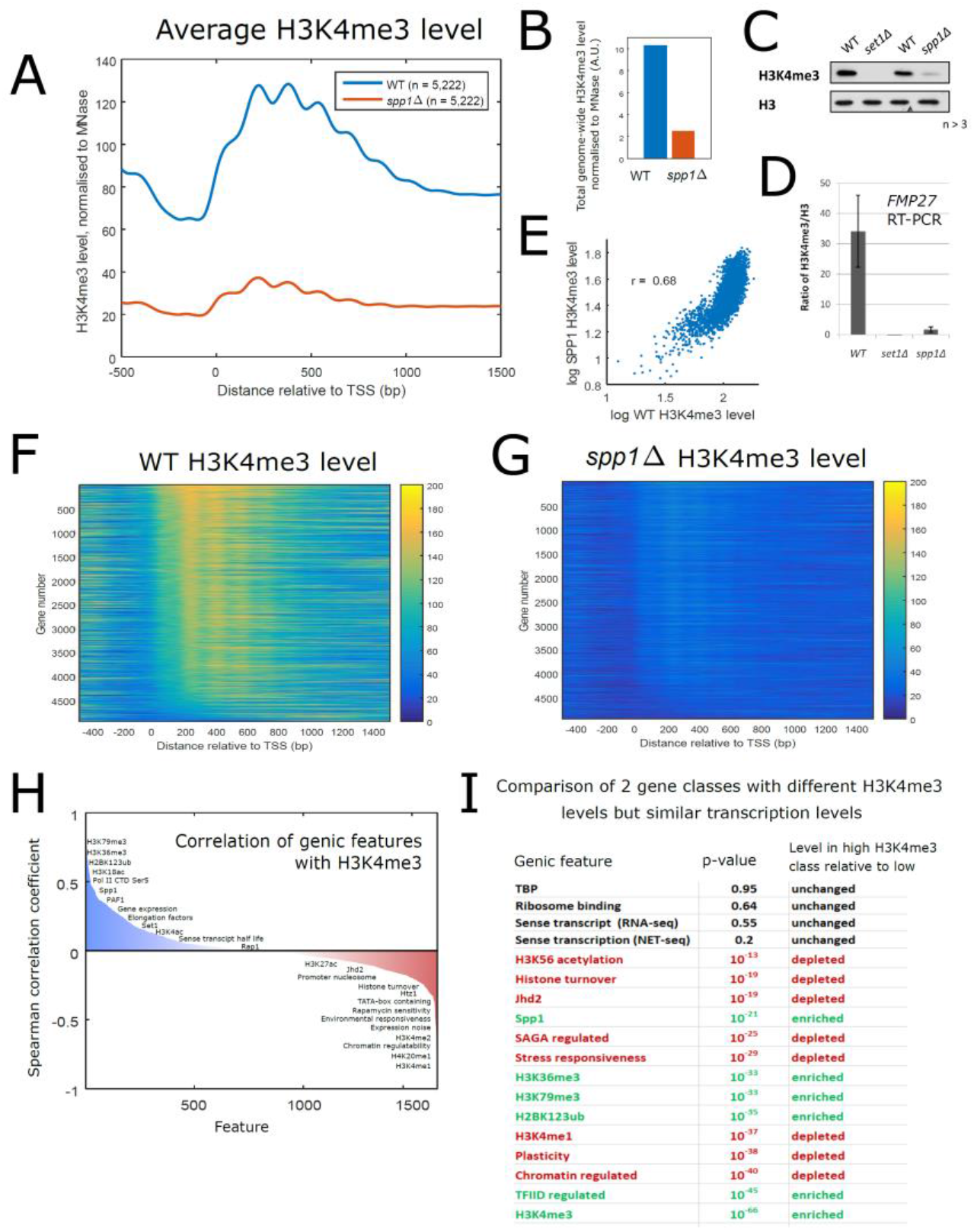
Deletion of *SPP1* massively ablates H3K4me3 levels genome wide. **(A)** The average H3K4me3 levels at 5,222 *S. cerevisiae* genes before and after deletion of *SPP1*, determined with ChIP-seq and spiking with *S. pombe* cells to allow for direct comparison between wild-type and *spp1Δ* strains. **(B)** Total genome-wide read count in wild-type and *spp1Δ* strains after normalisation by the spike-in control. **(C)** Northern blot showing total H3K4me3 levels in wild-type, *set1Δ* and *spp1Δ* strains, with H3 levels shown as a loading control. **(D)** HK4me3 levels at the *FMP27* gene determined by ChIP real time PCR, in wild-type, *set1Δ* and *spp1Δ* strains. **(E)** Scatterplot comparing H3K4me3 levels at 5,222 genes. Shown is the Spearman correlation coefficient. **(F-G)** Levels of H3K4me3 at 5,222 genes in wild-type and *spp1Δ* strains, displayed as heat maps. Genes were sorted according to the level of H3K4me3 in the region 300bp downstream of the TSS. **(H)** Spearman correlation coefficients of >1,500 gene features compared with H3K4me3, ranked by coefficient value. Features of note are displayed. **(I)** Comparison of 2 gene classes selected so they had very different levels of H3K4me3 but moderate levels of transcription that were not significantly different. Classes were compared for levels of the same genic features as in H, using the Wilcoxon rank sum test. Shown are selected features with the accompanying p-value and whether or not they were depleted or enriched in the high H3K4me3 class compared to the low class. Features are ranked according to p-value.

There are numerous known associations between H3K4me3 and other genomic features and gene parameters, most famously the level of gene activity (Santos-Rosa et al., 2002). To gain some insight into how these many associations rank against one another, we collated a wide range of genome-wide data-sets and assessed how they each correlated with the levels of H3K4me3 at the 5’ region of genes. Included in this list were numerous chromatin modifications and modifiers, transcription factors, elongation factors, as well as gene parameters like transcription level, transcript level, translation level, gene expression noise and the response of expression to mutations and environmental changes (sometimes referred to as *plasticity)* (Tirosh and Barkai, 2008). This amounted to over 1,500 different measurements. Each feature was correlated with H3K4me3 and a Spearman correlation coefficient obtained (Fig 1H).

Reassuringly, our H3K4me3 data correlated more strongly with other existing H3K4me3 datasets than any other feature (r=0.6-0.8). This was followed by the level of H3K79me3 (r=0.6), then H2BK123 ubiquitination (r=0.59), and then H3K36me3 (r=0.56), all measured across the body of the gene. Following this was the level of RNAPII CTD phosphorylation of Ser5 at the 5’ end (r=0.48), and Spp1 itself (r=0.48), also at the 5’ end. The various measures of gene expression ranked below these, with levels of transcription determined by NET-seq ranking the highest (r=0.44). The relationships observed were largely repeated when using an alternative source of genome-wide H3K4me3 data (Kirmizis et al., 2009), and were largely inverted when correlating features with H3K4me2 instead of H3K4me3 (Fig S1)((Kirmizis et al., 2007), as one might expect given the mutually exclusive nature of these marks.

Looking at the features that correlated negatively with H3K4me3, we found that levels of H3K4me1 at the 5’ end correlated most strongly (r=-0.61), followed by H4K20me (r=-0.56) and H3R2me2a (r=-0.54). Other strong, negative correlations included sensitivity to various different mutations (the largest being r=-0.50) as well as responsiveness to changing environmental conditions (r=-0.32), the presence of a TATA-box (r=-0.27) and expression noise (r=-0.32). This final association fits with a previously identified link between H3K4me3 and transcriptional consistency in human cells (Benayoun et al., 2014).

We were interested in determining to what extent these relationships could simply be explained by the correlation between H3K4me3 and gene expression. To this end we defined gene groups, of 1100 and 160 genes respectively, which had levels of transcription (determined by NET-seq) that were not significantly different (p = 0.2, Wilcoxon rank sum test) but very different levels of H3K4me3 (Fig S2), with one class having very high levels of H3K4me3 (greater than the 50^th^ percentile) and the other class having very low levels (less than the 15^th^ percentile). This allowed us to assess what features remained distinct between the two gene classes despite the similar levels of transcription (Fig 1I).

Strikingly, despite there being no statistically significant difference in the levels of transcript, transcription, TBP or ribosome binding, we otherwise observed very similar trends to those noted above (Fig 1H), with the high H3K4me3 group having high levels of H3K36me3/H3K79me3/H2BK123ub and being predominantly TFIID regulated, and the low H3K4me3 group being highly plastic, frequently containing a TATA-box, and having higher levels of histone turnover and H3 acetylation. This lower H3K4me3 class were also more responsive to stress and to mutations in various chromatin regulators, and had higher cell-to-cell variation in protein levels. This demonstrates that even when comparing groups with similar levels of gene-expression, differences in H3K4me3 are related to differences in a number of *dynamic* features of genes, both in terms of chromatin and gene regulation.

Taken together, our broad analysis indicates that genes with high levels of H3K4me3 tend to be highly expressed, TATA-less genes with a low variability in expression levels from cell-to-cell and condition-to-condition, and are also likely to be marked by H3K36me3, H3K79me3, H2BK123 ubquitination and RNAPII CTD serine 5 phosphorylation.

### Genome-wide reduction of H3K4me3 does not alter transcription or transcript patterns

The crosstalk between H2B ubiquitination, H3K79me3 and H3K4me3 is well-established (Zhang et al., 2015) and causal links identified, however as we have discussed before (Howe et al., 2017), there is a lack of convincing evidence to support a role for H3K4me3 in actually instructing transcription, despite the correlations described above. To assess whether H3K4me3 might instruct transcription, we measured genome-wide levels of both transcript and transcription in an *spp1Δ* strain, by RNA-seq and NET-seq respectively. NET-seq is a modification of RNA-seq in which RNAPII is first immunoprecipitated, and the nascent RNA purified prior to sequencing, such that the level of *transcription* is measured rather than steady-state transcript level (Churchman and Weissman, 2011). Our RNA-seq data was obtained in duplicate using an *S. pombe* spike-in protocol, allowing for better comparisons between wild-type and the *spp1Δ* strain.

Strikingly, we found there were no significantly enriched or depleted genes when comparing our RNA-seq data between *spp1Δ* and wild-type (Fig 2A), despite the massive reduction in levels of H3K4me3. Grouping genes according to various different classifications did not reveal any unique behaviours (Fig 2B). This initial observation strongly argues against any role of H3K4me3 in directing transcription. What is more, when assessing the distribution of transcript levels at genes, we observed only a very small, though significant reduction in *spp1Δ* compared to WT (Fig 2C).

**Figure 2.**
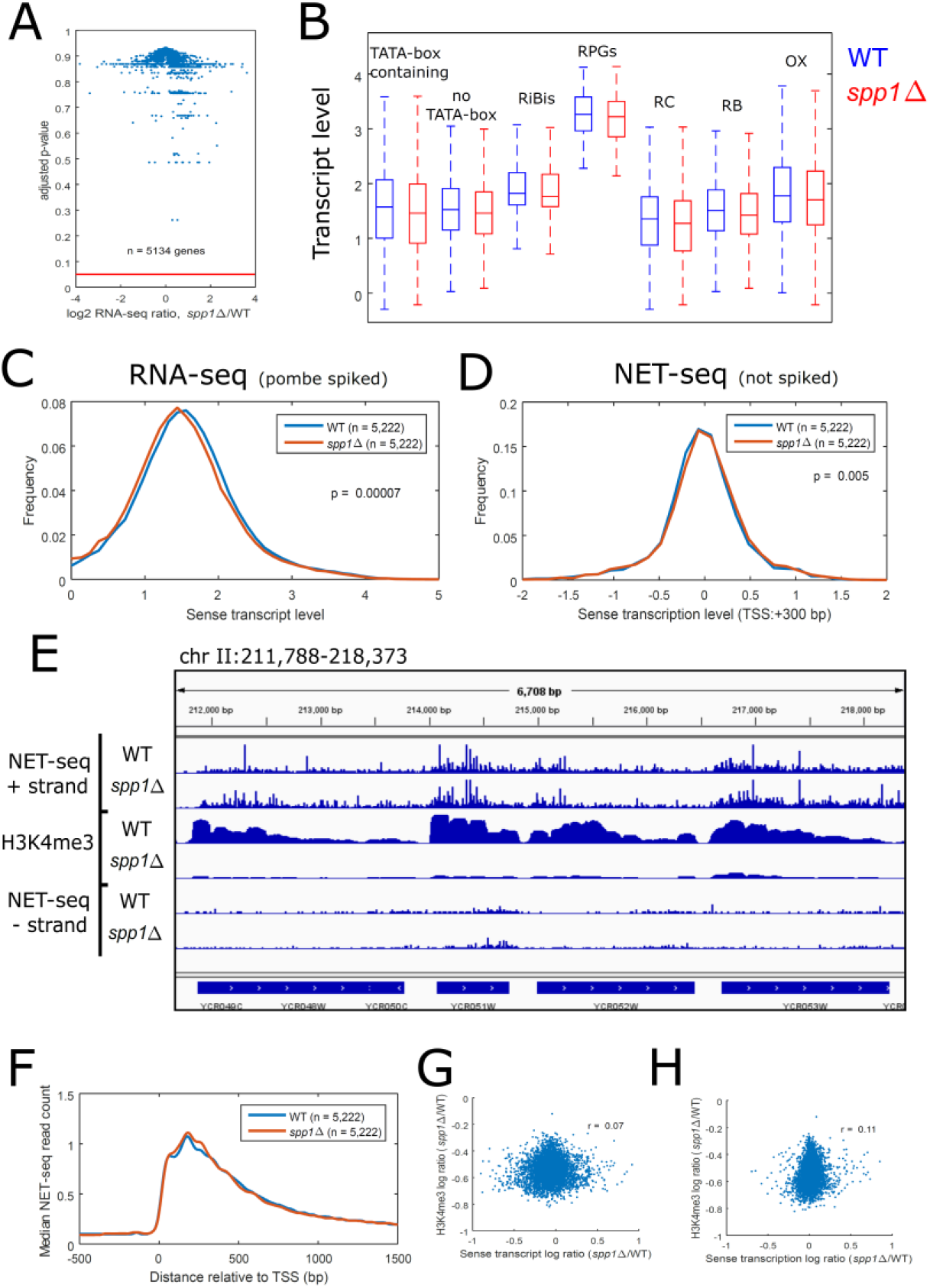
Ablation of H3K4me3 has little effect on genome wide transcript and transcription levels. **(A)** Volcano plot of transcript levels for 5,134 *S. cerevisiae* genes determined by RNA-seq, in both wild-type and *spp1Δ* strains. Red line indicates an adjusted p-value of 0.05. **(B)** Box plot showing transcript levels, determined by RNA-seq, at a number of different subclasses of genes, in both wild-type and *spp1Δ* strains. **(C)** Distribution of transcript levels of 5,222 genes in both wild-type and *spp1Δ* strains, determined by RNA-seq. **(D)** Distribution of transcription levels of 5,222 genes in both wild-type and *spp1Δ* strains, determined by NET-seq. **(E)** Snapshot of 4 genes on chromosome III, showing levels of transcription and H3K4me3 (determined by NET-seq and ChIP-seq respectively in both wild-type and *spp1Δ* strains. **(F)** Average number of NET-seq reads across a gene in both wild-type and *spp1Δ* strains. **(G)** Scatterplot comparing the ratio of transcript levels (determined by RNA-seq) at genes in wild-type and *spp1Δ* strains with the ratio of H3K4me3 in the same strains. Shown is the Spearman correlation coefficient. **(H)** Scatterplot comparing the ratio of transcription levels (determined by NET-seq) at genes in wild-type and *spp1Δ* strains with the ratio of H3K4me3 in the same strains. Shown is the Spearman correlation coefficient.

To assess whether H3K4me3 might play a role in transcription elongation, we obtained genome-wide NET-seq data in wild-type and *spp1Δ* strains. Churchman and Weissman (2011) previously demonstrated that deletion of the general elongation factor Dst1 (TFIIS) resulted in a substantial change to the average NET-seq profile, with the signal skewed much more towards the 5’ end of the gene. However, following deletion of Spp1, we observed no real change in the distribution of NET-seq reads amongst genes (Fig 2D-E), or in the average NET-seq metagene profile (Fig 2F), suggesting H3K4me3 plays no role in modulating transcription elongation and/or pausing.

There was some variation in how much H3K4me3 was lost from different genes following Spp1 deletion. If transcription is dependent on H3K4me3, then we might expect the genes which lose the most H3K4me3 to also lose the most transcription. However we observed only very weak correlations between the ratio of H3K4me3 and the ratio of transcript (r= 0.07; Fig 2G), and the ratio of H3K4me3 and the ratio of transcription (r= 0.11; Fig 2H). This fits with the observation that there is no statistically significant change in gene transcript level following Spp1 deletion (Fig 2A).

It has been shown that H3K4 methylation can modulate recruitment of Nrd1, a factor which in turn mediates targeting of transcripts to the exosome for degradation (Terzi et al., 2011). To this end we sought to measure whether or not H3K4me3 might affect transcript stability. To achieve this, we utilised an anchor-away system (Haruki et al., 2008) in which RNAPII itself was depleted from the nucleus. RNA-seq was performed on aliquots taken at six time-points from 0 to 120 minutes after RNAPII depletion, using *S. pombe* spike-ins in order to compare across time-points. By fitting an exponential curve to the data we were able to calculate half-lives for every gene, some examples of which are shown in Fig 3A. Our half-lives showed a modest agreement with other published transcript half-life estimates (Fig S3)(Geisberg et al., 2014; Miller et al., 2011), though it should be noted that these different estimates are themselves at best only modestly correlated with one another (Fig S3).

**Figure 3.**
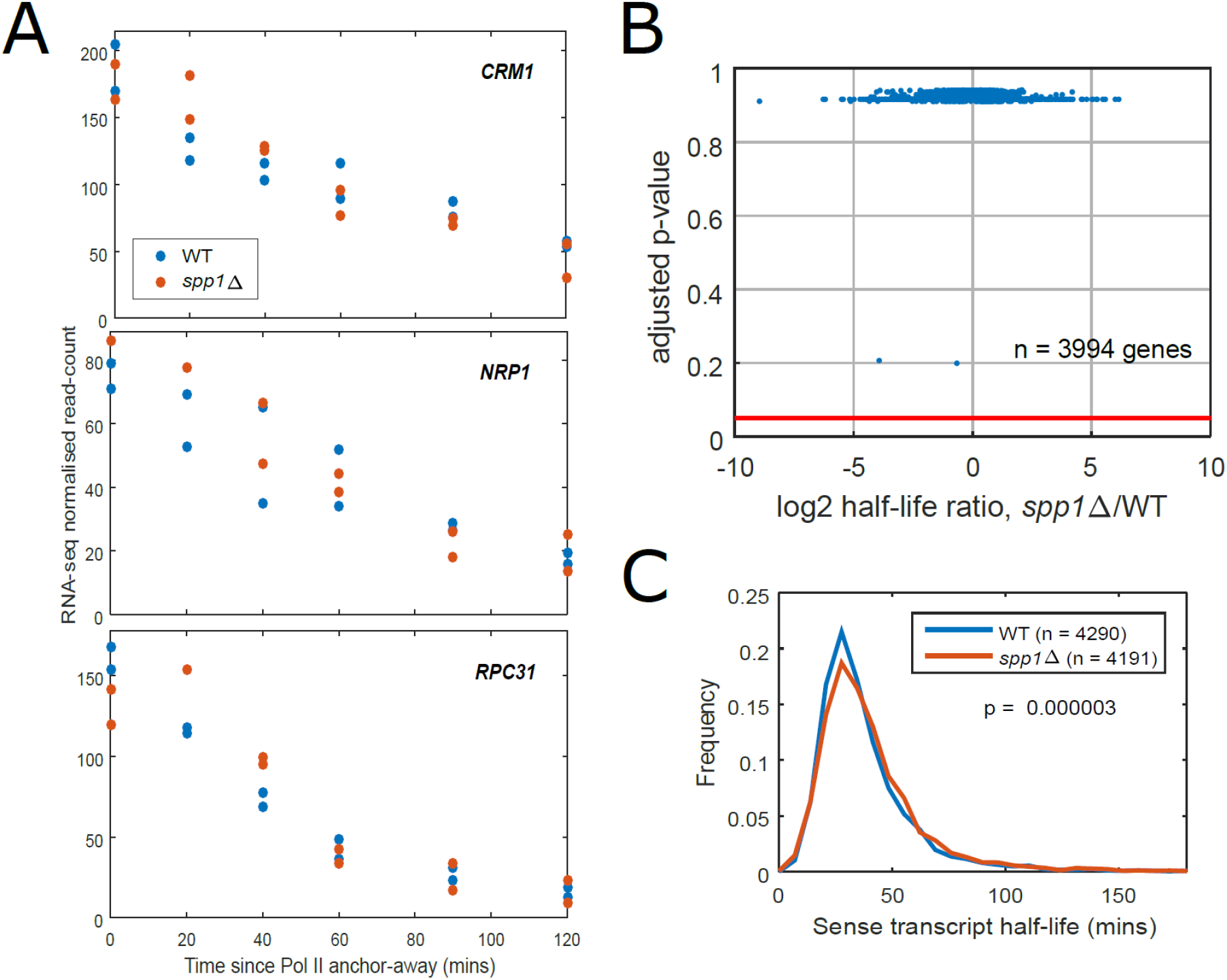
Transcript stability is not dependent on H3K4me3. **(A)** Loss of transcript at three different genes following depletion of RNAPII from the nucleus via an anchor away system, measured by RNA-seq. Transcript half-lives were determined for every gene using this data, in both wild-type and *spp1Δ* strains. **(B)** Volcano plot of half-lives for 3,994 genes, in both wild-type and *spp1Δ* strains. Red line indicates an adjusted p-value of 0.05. **(C)** Distribution of transcript half-lives in both wild-type and *spp1Δ* strains.

We observed no significant change in half-lives for any gene following Spp1 deletion (Fig 3B), and only a very slight though significant increase in average half-life when looking at overall half-life distributions (Fig 3C). Taken all together, the above results demonstrate that loss of H3K4me3 has no effect on mRNA biosynthesis, neither at the level of transcription, steady-state transcript level, or transcript degradation.

### Reduction of H3K4me3 does not alter transcription dynamics, translation or gene regulation

While it does not appear to alter overall levels of transcript or transcription, we were interested to assess whether H3K4me3 might modulate gene *dynamics*, as suggested by the above associations with expression noise. To this end, we utilized our previously-developed mathematical model (Brown et al., 2018), which describes how the underlying dynamics of transcription give rise to the populations of nuclear and cytoplasmic transcripts observable with RNA-FISH data. We have previously used this model to demonstrate how non-coding antisense transcription alters transcription and transcript processing events without changing overall levels of steady-state coding transcripts (Brown et al., 2018).

Building on existing models of transcription (Choubey et al., 2015; Raj et al., 2006; Zenklusen et al., 2008), our stochastic model allows us to discriminate between transcriptional events at the sense promoter, the nucleus and the cytoplasm. Briefly, our model describes a gene promoter switching stochastically between an active and inactive state, where initiation occurs only in the active state. Following this, the time taken for RNAPII to advance along the DNA and for the transcript to be transported from the nucleus is modelled as a series of stochastic jumps. Finally, transcripts are assumed to decay with a constant half-life. Our model allows us to derive initiation rates and ‘nuclear processing rates’, which is an umbrella term for the processes of elongation and nuclear export. When compared with experimental data, the model allows us to determine which parameters of transcript production and processing change between wild-type and *spp1Δ* strains.

We performed RNA-FISH on twelve different genes in both wild-type and *spp1Δ* strains. We deliberately selected genes with varying levels of H3K4me3, and which showed a substantial reduction of H3K4me3 following *SPP1* deletion. By simulation we obtained the parameters that best described the observed data, and used these to determine the mean initiation rates and nuclear processing rates for all twelve genes in both strains.

Our obtained initiation and nuclear processing rates showed a moderate correlation with the level of H3K4me3 at the same genes (Fig S4, r = 0.2 and 0.3 respectively, Spearman correlation coefficient). However, comparison of rates between wild-type and *spp1Δ* strains showed very little deviation, for both initiation rates and nuclear processing rates (Fig 4A). Any small changes observed were not consistent across the twelve genes, leading us to conclude that loss of H3K4me3 does *not* lead to a change in transcription dynamics, not at the level of initiation, elongation or nuclear export.

**Figure 4.**
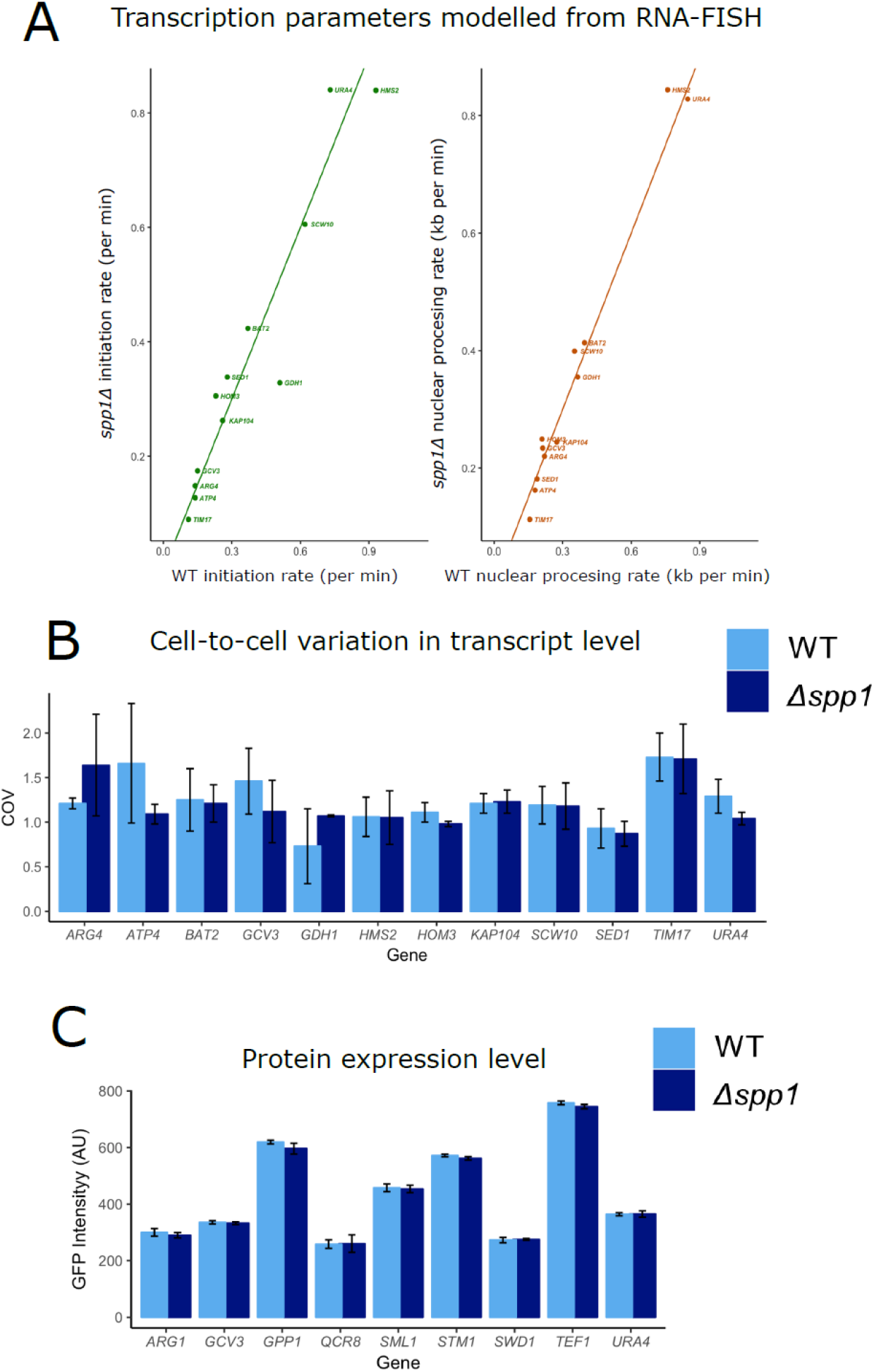
Ablation of H3K4me3 does not alter gene expression parameters. (A) RNA-FISH data for 12 genes was obtained in both wild-type and *spp1Δ* strains, and used to estimate the initiation rates and nuclear processing rates at these genes (see text). Here, transcription parameters for wild-type are plotted against those for *spp1Δ*. Any great difference would be observable as a deviation from the straight line (B) Cell-to-cell variation in transcript level for 12 genes in both wild-type and *spp1Δ* strains, determined with RNA-FISH, measured as the coefficient of variation in average cytoplasmic signal across several hundred cells per gene per strain. (C) Levels of protein expression for 8 GFP-tagged genes, determined by FACS in both wild-type and *spp1Δ* strains.

Given the association between H3K4me3 and expression noise shown in Fig 1H, and described in Benayoun et al. (2014), we also assessed whether the cell-to-cell variation of transcript level was altered by deletion of Spp1. However, we saw only small changes in the coefficient of variation (COV), which were not consistent across the twelve genes studied (Fig 4B). This demonstrates that the low cell-to-cell variation found at genes with high H3K4me3 is not itself maintained by the Spp1-deposited H3K4me3.

Next, we wished to assess whether H3K4me3 might modulate gene expression at the level of translation. We used FACS to measure the expression levels of nine of the twelve genes investigated above, by tagging them with GFP. Consistent with the effects on transcription, we saw only very small, inconsistent changes in average protein level across these twelve genes, following deletion of Spp1 (Fig 4C).

So far, our analysis has focussed entirely on yeast existing in steady-state conditions. It is possible however that H3K4me3 might affect the manner in which genes respond to environmental changes, as suggested by the above, negative association with environmental responsiveness (Fig 1H). To this end, we obtained NET-seq data in *S. cerevisiae* grown in glucose then induced with galactose, in both wild-type and *spp1Δ* strains. We hypothesised that if H3K4me3 represses a gene’s ability to respond to environmental changes, then we would observe a greater ‘inducibility’ at genes as a consequence of a loss of H3K4me3.

Galactose has a profound effect on the genome-wide patterns of transcription in yeast. We identified a low correlation between the number of NET-seq reads at genes in glucose and galactose (r = 0.44, spearman correlation coefficient, Fig 5A). 277 genes showed a greater fold increase in transcription than the well-studied galactose-inducible gene *GAL1*. However, wild-type and *spp1Δ* strains behaved highly similarly in their response to galactose induction (Fig B-D). NET-seq data was extremely well correlated between the two strains, both after 15 minutes and 60 minutes in galactose (r = 0.99 and 0. 98 respectively, Fig 5B). Furthermore, the change in transcription levels from glucose to 60 minutes in galactose was highly correlated between the two strains (r = 0.9, Fig 5C), demonstrating that genes respond the same way to galactose irrespective of the presence or absence of H3K4me3. Of the 200 genes most enriched in the wild-type strain following 60 minutes in galactose, 170 were present within the 200 most enriched genes in the *spp1Δ* strain (the remaining 20 were within the top 300). Comparing the distribution of changes showed that *spp1Δ* showed no particular preference for overall enrichment or depletion of transcription level compared to wild-type (Fig 5D). Taken together, our results demonstrate that H3K4me3 plays no role in modulating the genomic response to galactose.

**Figure 5.**
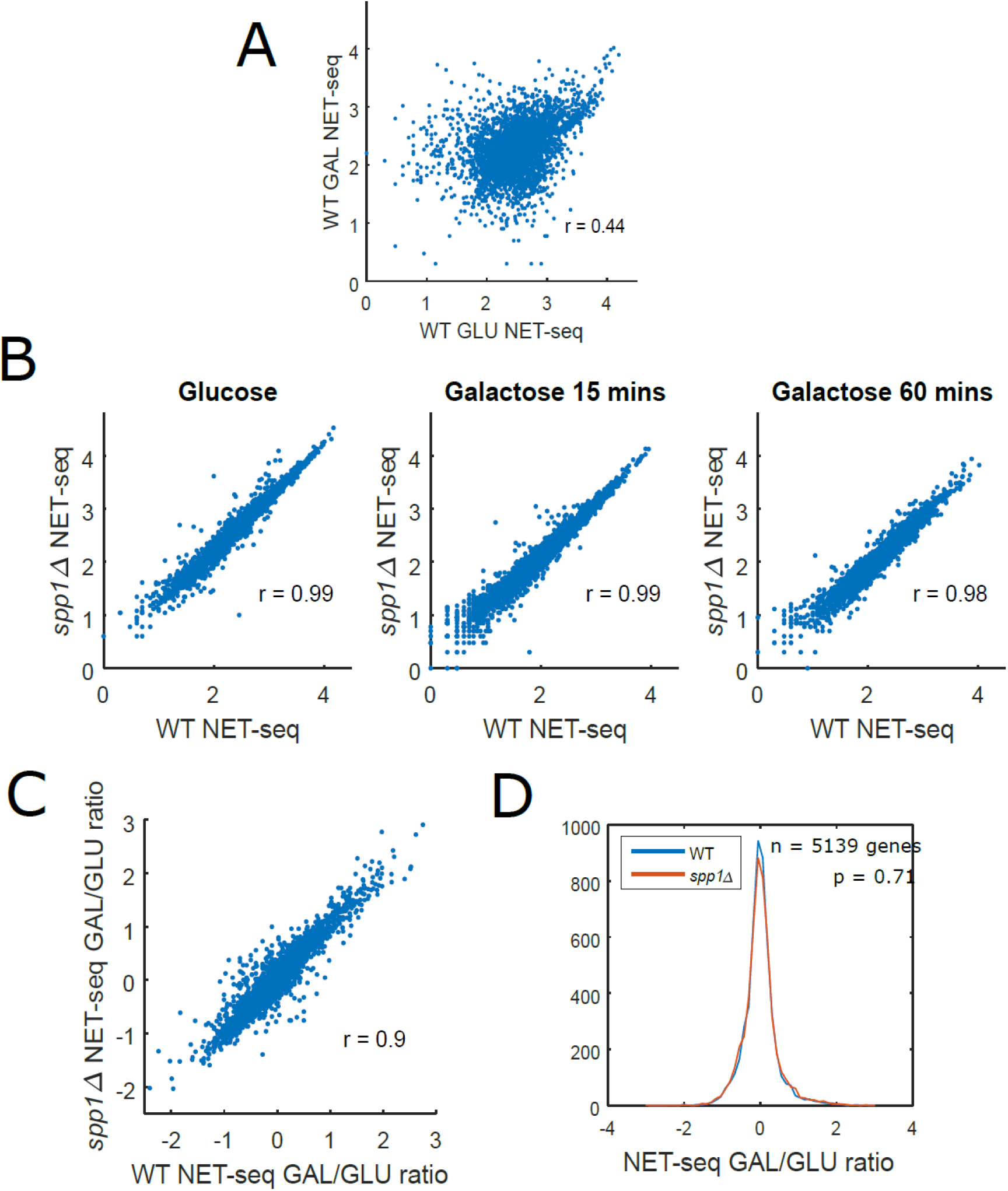
The large genomic transcription response to galactose is near identical following ablation of H3K4me3. **(A)** Scatter plot of transcription levels of 5,222 genes in glucose versus one hour in galactose, determined by NET-seq. Shown is the Spearman correlation coefficient. **(B)** Scatterplots of transcription levels of 5,222 genes in wild-type versus *spp1Δ*, in glucose, 15 minutes galactose and 60 minutes galactose, determined by NET-seq. Shown are the Spearman correlation coefficients. **(C)** Scatter plots of the changes in transcription level for 5,222 genes following one hour in galactose, in wild-type versus *spp1Δ*. Shown is the Spearman correlation coefficient. **(D)** Distribution of the ratio of transcription levels before and after one hour galactose induction, in both wild-type and *spp1Δ* strains.

### Changes in genome-wide transcription patterns do not alter the maintenance or deposition of H3K4me3

We hypothesised therefore that H3K4me3 levels might be dependent on transcription, and not vice versa. To this end, we measured the genome-wide levels of H3K4me3 following the depletion of RNAPII from the nucleus via an anchor away system, at 0, 20, 40 and 60 minutes following the addition of Rapamycin. Using an FRB-GFP tagged Rpb1 (the largest subunit of RNAPII), we observed that the polymerase is spread out entirely in the cytoplasm by 60 minutes (Fig 6A). This loss of RNAPII resulted in a dramatic reduction in global transcript levels (Fig 6B).

**Figure 6.**
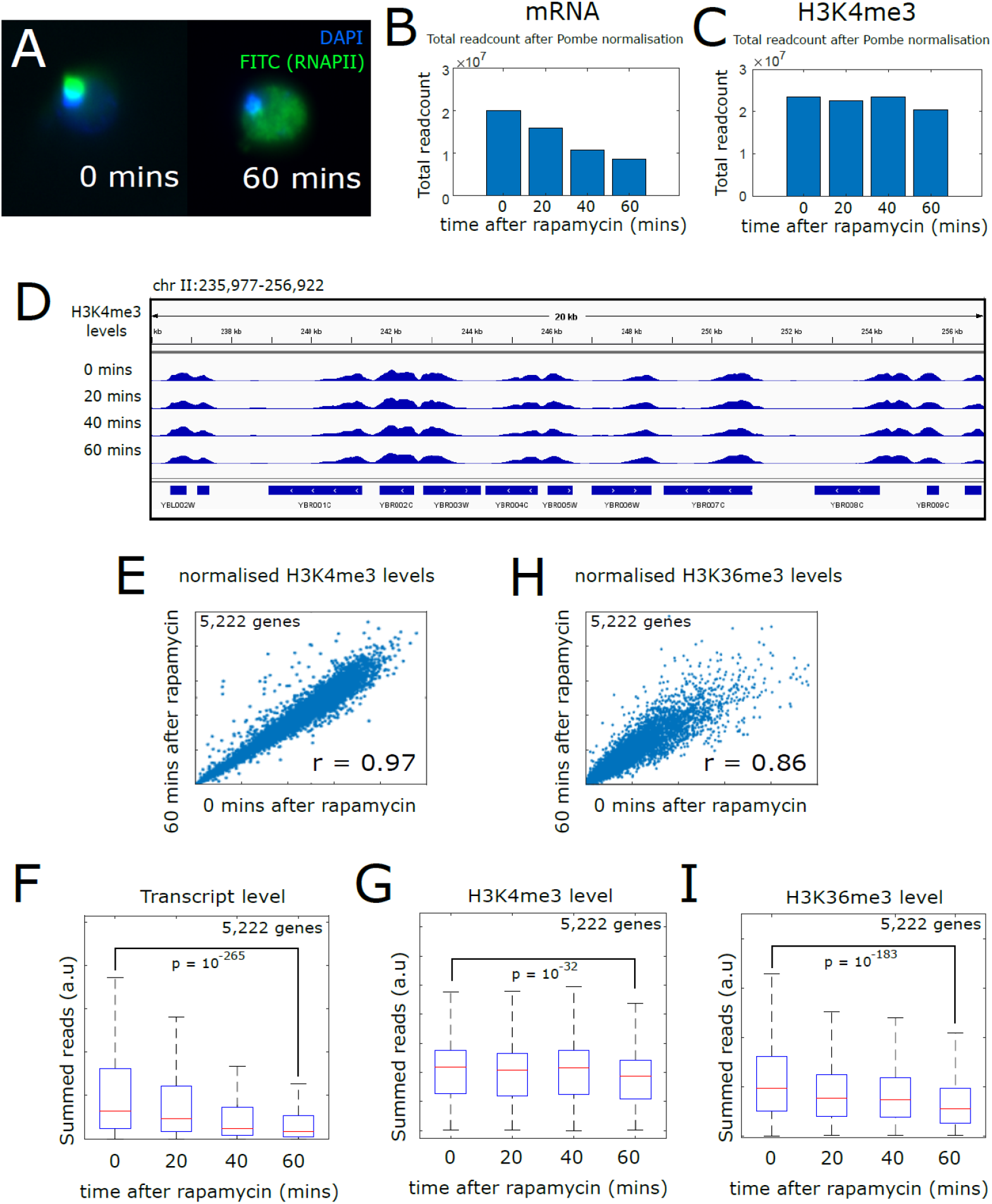
Depletion of RNAPII from the nucleus alters some chromatin but not H3K4me3. **(A)** Fluorescence microscopy of cells with GFP-tagged RNAPII, 0 and 60 minutes after the anchor away system was induced by the addition of rapamycin. **(B-C)** Total genome-wide read counts for both mRNA (determined by RNA-seq) and H3K4me3 (determined by ChIP-seq) following addition of rapamycin and depletion of RNAPII from the nucleus. **(D)** Snapshot of 12 genes on chromosome II, showing levels of H3K4me3 determined by ChIP-seq up to 60mins after the addition of rapamycin and the depletion of RNAPII from the nucleus. **(E-H)** Scatterplot comparing H3K4me3 and H3K36me3 levels at 5,222 genes, at 0 versus 60 minutes following the addition of rapamycin. Shown is the Spearman correlation coefficient. **(F-I)** Boxplots showing levels of mRNA transcript, H3K4me3 and H3K36me3 at 5,222 genes following the addition of rapamycin and depletion of RNAP1mII from the nucleus. Displayed p-values were obtained by a Wilcoxon rank sum test comparing genes at 0 mins with genes at 60 mins.

Strikingly, depleting RNAPII from the nucleus has little effect on H3K4me3 (Fig 6C-D). By spiking our ChIP-seq experiments with *S. pombe* cells we were able to compare total levels of H3K4me3 between different time points, and observed little difference (Fig 6C). Furthermore, the levels of H3K4me3 at individual genes was highly similar across time points (Fig 6D-E, spearman correlation coefficient = 0.97). The level of H3K4me3 in the 5’ gene region varied little following RNAPII depletion compared to the dramatic reduction in transcript level (Fig 6F-G). We therefore conclude that both the genome-wide patterns and global levels of H3K4me3 do not change following a loss of transcription.

For comparison, we also assessed how another modification, H3K36me3, changed following depletion of RNAPII from the nucleus. Global patterns of H3K36me3 were not quite so well-correlated across time points as H3K4me3 (Fig. 6H, spearman correlation coefficient = 0.86), but perhaps more strikingly, showed a much more noticeable reduction at genes (Fig. 6I). This suggests that histone modifications are not always slow to respond to the cessation of transcription, and that while some chromatin features do respond to changes in transcription, H3K4me3 is particularly insensitive.

Weiner et al. (2015) have previously studied the response of a wide set of histone modifications to diamide stress in budding yeast. We reanalysed this data, focussing on H3K4me3 levels and how their changes are related to changes in transcription levels, as assessed by genome-wide RNAPII ChIP (Kim et al., 2010). We find that despite broader changes in transcription, the genome-wide pattern of H3K4me3 changed very little in response to diamide (Fig 7A-B). The correlation between RNAPII levels at genes at 0 and 60 minutes after diamide was much lower than the same correlation for H3K4me3 (spearman correlation coefficients = 0.69 vs 0.97), suggesting H3K4me3 levels are more resistant to environmental stimuli than transcription. Furthermore, what little variation was present showed no correlation with the variation in transcription (r = 0.01). This fits with our above observation that depletion of RNAPII from the nucleus has little effect upon H3K4me3. Looking at two other modifications, we found that global patterns of H3K36me3 were also relatively unchanged after 60 minutes (r = 0.95; Fig. 7A), while acetylation of H3K14 showed larger changes, closer to those observed for RNAPII levels (r = 0.78; Fig. 7A). When we isolated the 250 genes that showed the greatest increase and decrease in RNAPII levels in response to diamide stress, and compared the H3K4me3 levels in the same groups, we observed no major changes over time (Fig 7B). H3K36me3 levels also showed little change after 60 minutes, while H3K14ac levels rose in those 250 genes at which RNAPII levels increased in diamide (Fig 76B).

**Figure 7.**
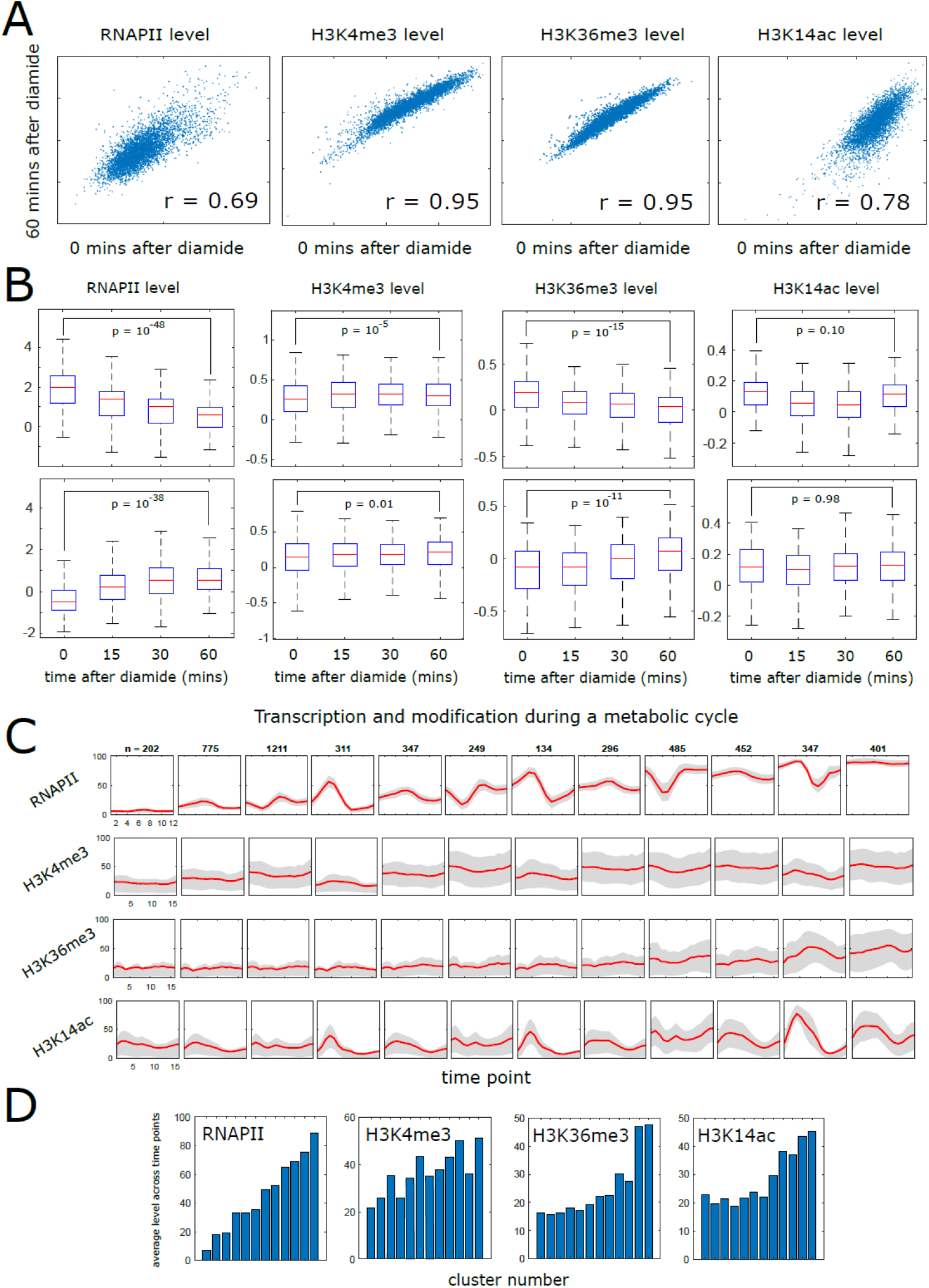
H3K4me3 changes little in response to changes in transcription. **(A)** Scatterplots comparing levels of different factors at genes 0 and 60 minutes following the addition of diamide; RNAPII, H3K4me3, H3K36me3 and H3K14ac. **(B)** Boxplots showing levels of the same factors following addition of diamide. Top: the 5% of genes which showed the biggest decrease in RNAPII following addition of diamide. Bottom: the 5% of genes which showed the biggest increase in RNAPII. **(C)** Levels of these same factors in twelve different subsets of genes during the yeast metabolic cycle, subdivided by k-means clustering according to the behaviour of transcription during a single metabolic cycle, measured by NET-seq. **(D)** Levels of RNAPII, H3K4me3, H3K36me3 and H3K14ac within each of these 12 clusters, averaged across all measured time points of a single metabolic cycle.

Budding yeast undergo cyclical changes of gene expression when grown under continuous, nutrient-limited conditions (Tu et al., 2005). We reanalysed the levels of H3K4me3 measured at 16 points within a single cycle (Kuang et al., 2014), to determine whether they change concurrently with changes in transcription. To achieve this we determined genome-wide transcription levels at 12 time points within a single cycle, using NET-seq (unpublished data). We used k-means clustering to divide genes according to how their sense transcription changes during the cycle, then assessed the levels of H3K4me3 in the same groups. Levels of both transcription and modification were expressed as a rank, i.e. the percentile the level of a gene occupied compared against all other genes across all time points. We observed that while transcription showed large changes over the cycle, H3K4me3 did not show large changes in these same groups (Fig 7C). This was entirely different to what was observed for other histone modifications such as H3K14ac, which showed large changes over time, with the same trends as observed for transcription (Fig 7C). The average level of H3K4me3 across all time points did show a weak association with average RNAPII levels (Fig 7D), however the fact that H3K4me3 did not vary across time points demonstrates that it does not change as transcription does within a dynamic system.

Taken together, our findings demonstrate that although H3K4me3 marks highly expressed, environmentally-insensitive genes, it neither requires transcription for its deposition, nor is it required for the maintenance of the genome-wide gene-expression program nor the transcriptional response to environmental changes.

## Discussion

Here, we have sought to elucidate the relationship between H3K4me3 and gene expression, assessing the effects of a loss of H3K4me3 at every level of gene expression, and in turn observing how H3K4me3 levels respond to changes in transcription. We find that the reduction of global levels of H3K4me3 to 25% does not significantly alter the transcript levels of any gene, with no change to initiation rate, transcript stability, translation level, cell-to-cell transcript variability or the distribution of elongating polymerase on the gene. Loss of RNAPII from the nucleus does not change the genome-wide pattern of H3K4me3 and has little effect on global levels, while changes in transcription in response to environmental stresses are not reflected in changes to H3K4me3. Our findings demonstrate that despite their well-established correlation, H3K4me3 is not an activator of transcription, nor does it respond to changes in transcription. This demands a change to the simplistic view represented by both models, whereby increased H3K4me3 is not predictive of increased transcription and vice versa.

Here we report that deletion of Spp1 does not result in the differential expression of any gene at the level of mRNA transcript, while we have previously reported that only 57 genes are upregulated and 23 downregulated at the level of transcription (measured by NET-seq)(Howe et al., 2017). This is in agreement with a mammalian study by Clouaire et al. (2012), who found that CFP1 deletion only weakly affected transcription in embryonic stem cells, and that the changes in H3K4me3 resulting from CFP1 deletion were not related to changes in RNAPII occupancy. By contrast, it has been shown that deletion of CFP1 in mouse oocytes had a larger effect, with 718 genes showing a statistically significant increase and 382 a decrease (Yu et al., 2017), while in embryonic stem cells 584 genes showed a significant increase and 1,108 genes a significant decrease following CFP1 deletion, from a set of 9,979 genes (Brown et al., 2017), although these changes, while significant, were generally small, and loss of CFP1 (and thus H3K4me3) did not consistently upregulate or downregulate. It should be noted that, unlike Spp1, CFP1 has additional binding modules that target the COMPASS complex to promoters and enhancers, and as such these effects may not be the result of a loss of H3K4me3 but rather from distinct effects mediated by DNA binding (Hu et al., 2017).

If H3K4me3 is not generally instructive for transcription, and transcription is not necessary to maintain H3K4me3, then it follows that some other factor must be responsible for the association between the two. H3K4me3 levels are high at transcriptionally active genes, particularly those with low levels of cell-to-cell variability (noise) and condition-to-condition variability (plasticity), i.e. those that might traditionally be referred to as constitutive or ‘house-keeping’ genes. Possible candidates for regulators of H3K4me3 could be proteins that bind to the same subset of H3K4me3-enriched, transcriptionally active yet environmentally unresponsive genes, and which also support transcription. TFIID would be such a candidate, as a frequent component of the PIC which has previously been shown to bind preferentially to the promoters of unregulated genes (Huisinga and Pugh, 2004).

It is also possible that the establishment of a PIC, prior to the recruitment of RNAPII, is sufficient to recruit H3K4me3 – i.e. the process of initiation up until the involvement of the polymerase – though this does not agree with our observation that genes activated by diamide stress to not show a similar increase in H3K4me3 level. A further possibility is that the hypothetical transcription activatory role of H3K4me3 is shared amongst other modifications, and that they function redundantly in this fashion. Acetylation of H3 is associated with the level of transcription, as is H3K36me3, H3K79me3, and H2B ubiquitination. It is also possible that H3K4me3 and H3K4me2 function redundantly – it has been shown that in the absence of H3K4me3, the peak of H3K4me2 is shifted to occupy the site on the gene where H3K4me3 would normally be present (Soares et al., 2017). However this is at odds with observations that the readers of H3K4me3 and H3K4me2 can be highly specific (Shi et al., 2006).

The presence of H3K4me3 predominantly at environmentally unresponsive genes, twinned with the observation that H3K4me3 changes very little in response to diamide stress, suggests the possibility that H3K4me3 levels are themselves environmentally unresponsive. There is a general lack of available data comparing genome-wide levels of H3K4me3 (and histone modifications generally) in distinct environments, and further investigation would help to elucidate whether H3K4me3 is generally unresponsive to large scale changes in the transcriptional program of the cell. Ng et al. (2003) have proposed that H3K4me3 might play the role of a transcriptional memory – a related hypothesis might be that H3K4me3 levels are instead a reflection of a gene’s ‘preferred’ or ‘basal’ transcription level in the absence of environmental cues, hence explaining why it is predominantly high at well-expressed, non-plastic genes, yet does not change when RNAPII is lost from the nucleus. We also find that H3K4me3 is not required to regulate the large transcriptional changes observed when yeast is shifted from a glucose to galactose-containing medium. This is similar to what has previously been observed in mouse embryonic stem cells, in which deletion of CFP1 has little effect on the transcriptional response to induction by doxorubicin, a DNA damaging agent which alters the expression of 1,264 genes (Clouaire et al., 2014).

We find that H3K4me3 levels are maintained in the absence of RNAPII, while H3K36me3 is not. H3K4me3 is predominantly localized to the +1 nucleosome, which has been shown to experience the highest levels of H3 turnover (Dion et al., 2007). This turnover would presumably replace trimethylated histones with un-trimethylated histones, resulting in a reduction. This begs the question of what exactly is maintaining the levels of this modification. As discussed above, one possibility for this is that some element of the transcription initiation machinery is responsible for Spp1 recruitment/H3K4me3 deposition, such as TFIID, which would explain the preference for H3K4me3 at the predominantly TFIID-regulated genes. The COMPASS complex can exist in a form lacking Spp1 (D’Urso et al., 2016), so it is entirely possible that some additional element would be required to remodel the complex to include Spp1. Set1 also contains an RNA-binding motif (RRM1)(Roguev et al., 2001), so it is possible the complex might be recruited by stable R-loops formed at the 5’ end of the gene that persist in the absence of RNAPII (Santos-Pereira and Aguilera, 2015), possibly established by non-coding antisense transcription.

Although the precise biological role of H3K4me3 is unclear, its importance is nevertheless apparent. In *S. cerevisiae*, loss of Spp1 (and therefore H3K4me3) is linked to poor chronological ageing (Cruz et al., 2018; Walter et al., 2014), while murine Cfp1–/– embryonic stem cells are incapable of differentiating (Carlone and Skalnik, 2001). In humans, chromosomal translocations in SET-containing genes are frequently associated with acute myeloid and lymphoid leukemias (Shilatifard, 2012). Furthermore, H3K4me3 has been shown to play a role in DNA repair (Faucher and Wellinger, 2010; Peña et al., 2008) and in the targeting of double-strand breaks during meiotic recombination (Acquaviva et al., 2013; Borde et al., 2009; Sommermeyer et al., 2013). Here we find that H3K4me3 is not necessary to maintain the genome-wide pattern of transcription, nor regulate the transcriptional response to environmental signals, nor is RNAPII necessary to maintain the pattern of H3K4me3. H3K4me3 is nevertheless a marker of a particular class of transcriptionally-active gene, namely those which show low variation in expression levels across a wide range of environmental conditions. Future research should focus on trying to ascertain why such genes should be marked in this way, particularly given that it doesn’t inform the transcription at these genes, to help elucidate its elusive biological function.

## Methods

### Yeast Culture and Genetic Manipulation

Yeast Culture and cloning procedures were performed as described previously (Brown et al., 2018).

### RNA-FISH and downstream analysis

RNA-FISH and modelling was performed as described previously (Brown et al., 2018).

### Anchor Away

Untagged WT and anchor-away strains were grown to log phase. Rapamycin was added to a final concentration of 1 ug/mL and samples for ChIP and Microscopy were taken every 20 min.

### ChIP-Seq

ChIPs were performed as described previously (Morillon et al., 2005) with the following exceptions: Before formaldehyde fixation, 25% (vol. at OD 0.6) *S.pombe* cells were added as a spike-in control. 10 uL of anti-H3K4me3 (Millipore, 05-745R) or anti-H3K36me3 (abcam, ab9050) were used for precipitation. Immunoprecipitated protein-DNA complexes were purified using protein-A dynabeads (Invitrogen). Libraries were prepared using the NEBNext Ultra II DNA Library Kit (NEB) and sequenced on an Illumina Nextseq 500 platform (Department of Zoology, Oxford).

### Microscopy

Microscopy was performed as described previously for RNA-FISH (Brown et al., 2018).

### FACS

2 mL mid-log phase cells were pelleted and resuspended in 4% paraformaldehyde/PBS. Cells were fixed for 45 min in the dark at room temperature with shaking at 70 rpm before resuspending in 400 uL PBS. For FACS measurements, cells were diluted 1:20 in PBS. 10,000 cells per sample were measured by FACS Calibur, set to a laser power of 15 mWatts.

### NET-Seq

NET-Seq was performed as described previously (Fischl et al., 2017).

### RNA-Seq

Before RNA purification, 25 % *S. pombe* cells were added (vol. at OD 0.6). Pelleted cells were resuspended in TES (10 mM Tris pH 7.5, 10 mM EDTA pH 8, 0.5 % SDS). RNA was then extracted with phenol:chloroform followed by ethanol precipitation. 500 ng of RNA was used for library preparation with the Lexogen 3’ end RNA-seq library prep kit. Libraries were sequenced on an Ion Torrent.

### Read alignment

Alignment of FASTQ read files was performed using Bowtie2 with standard settings. Where *S.pombe* spike-ins were used, reads were first aligned to the *S.pombe* 2007 build (ASM294v2) to determine the total number of aligned reads that would be used in the subsequent normalization. Reads that failed to align to *S.pombe* were then aligned to the *S.cerevisiae* sacCer3 build. Genome coverage files were generated with Bedtools and subsequent analysis performed using bespoke codes written in R and MATLAB.

### Bioinformatic analysis

Sense transcription start and end sites were determined as described previously (Brown et al., 2018), using transcript isoform sequencing (TIF-seq) data from Pelechano et al. (2013). Levels of nascent transcription, H3K4me3 and H3K14ac from all sources were determined as the average signal in a 300bp window placed immediately downstream of the TSS. Levels of H3K36me3 were determined as the average levels between the TSS and the transcription end site. Levels of transcript determined by 3’ end RNA-seq were obtained by measuring the average signal in a 300bp window placed immediately upstream of the transcription end site. TATA-box containing genes were those defined previously by Basehoar et al. (2004), RiBi and RPG classifications were obtained from the Saccharomyces Genome Database, while RC, RB and OX phase genes were those defined by Tu et al. (2005). All p-values were determined using the Wilcoxon rank sum test. Adjusted p-values were calculated using the Bonferroni Correction.

### Collated compendium of genome-wide factors

The details and sources of the list of factors features in Fig. 1H is available on request.

### Data availability

Raw data and processed read tables are available through the Gene Expression Omnibus (https://www.ncbi.nlm.nih.gov/geo/), under the SuperSeries GSE133838.

## Acknowledgements

We thank the J.M. laboratory for critical discussions, Anitha Nair for excellent technical support and Ilan Davis and Micron Oxford for microscopy support. This work was supported by: The Wellcome Trust (WT089156MA to J.M.); the BBSRC (BB/P00296X/1 to J.M.); the Leverhulme Trust (RPG-2016-405 to J.M.); a Wellcome Trust Strategic Award (091911) supporting advanced microscopy at Micron Oxford (http://micronoxford.com); an EPSRC studentship (EP/F500394/1 to T.B) and a Royal Society University Research Fellowship (UF120327 to A.A.).

## Author Contributions

Project conception: JM, SCM and FSH; NET-Seq: HF and SX; RNA-Seq, Northern blot and RT-PCR: FSH; ChIP-seq and anchor away: FSH and PL; FISH: MW; FISH downstream analysis: AA; mathematical modelling: TB and PL; FACS and strain construction: PL, WK and JRR; bionformatics: SCM. All authors were involved in interpreting the data. JM and SCM wrote the paper with input from PL.

---

**Figure S1.**
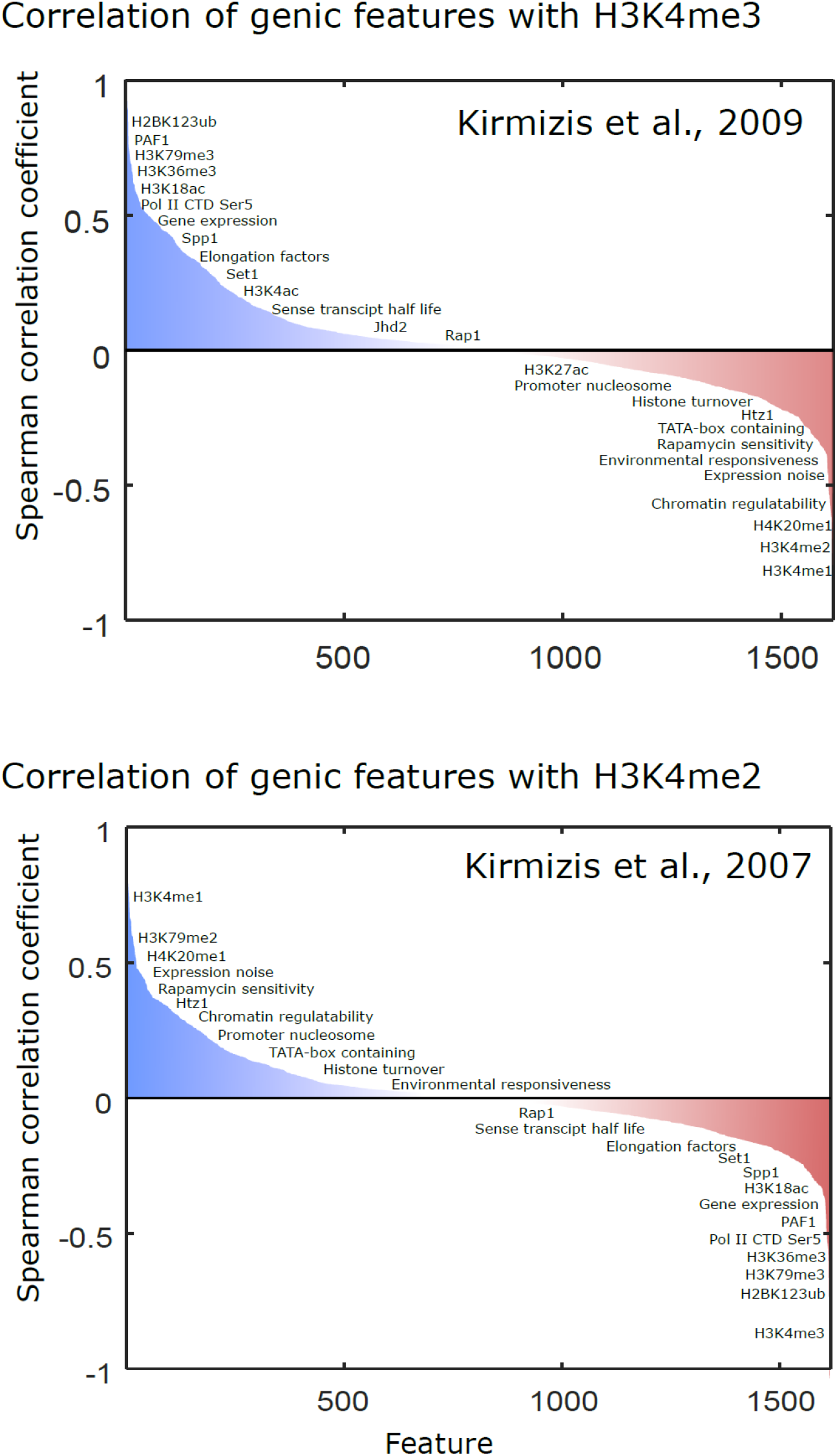
Correlation of >1,500 genic features with H3K4me3 and H3K4me2. Shown is the distribution of Spearman correlation coefficients for over 1,500 features, as summarised in the main text, for publically available genome-wide levels of H3K4me3 (top) and H3K4me2 (bottom). Certain features have been indicated.

**Figure S2.**
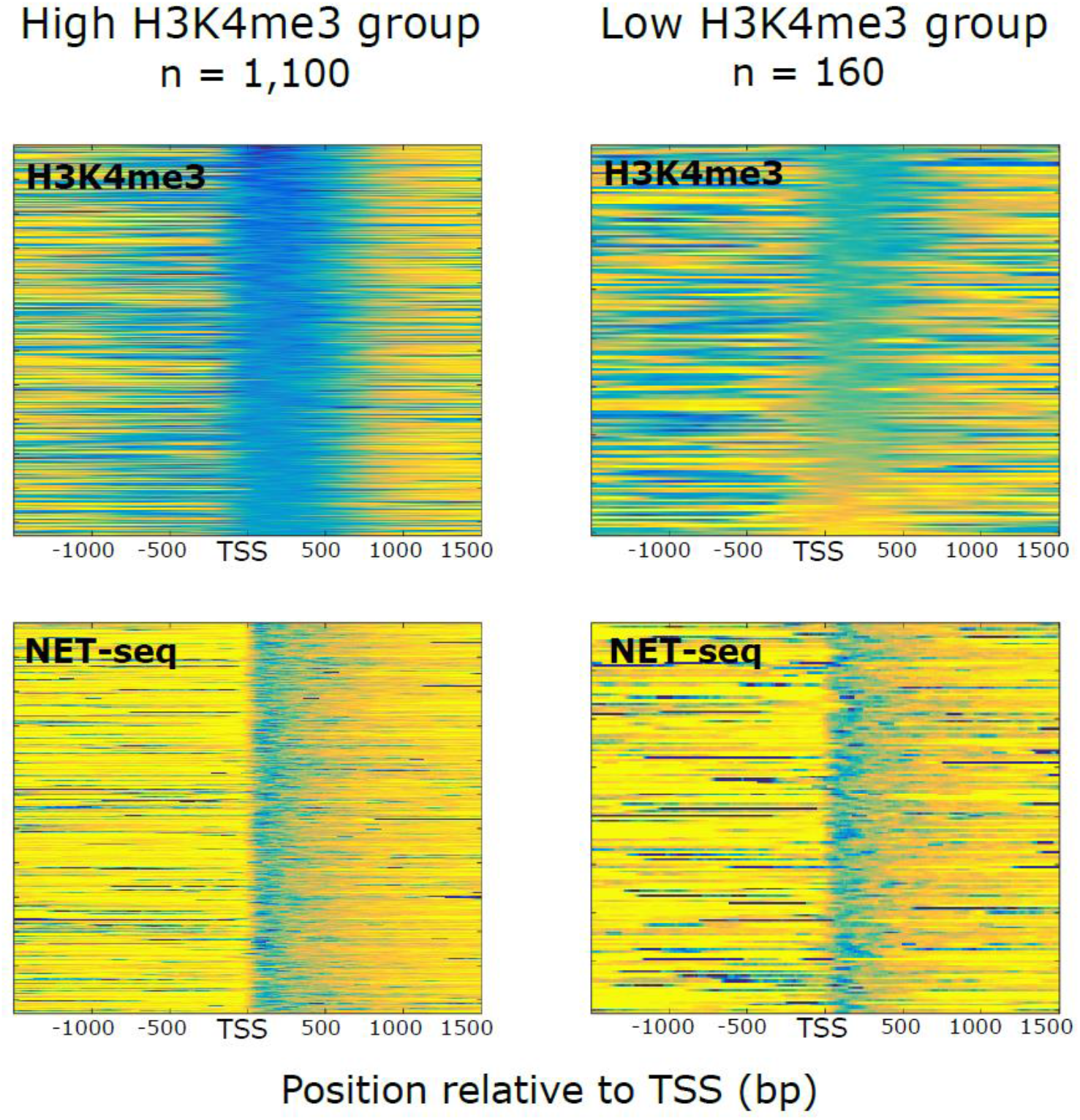
Two gene groups with similar transcription levels but different H3K4me3 levels. Two gene groups were isolated such that they had very similar levels of transcription but either high levels or low levels of H3K4me3, to demonstrate that pronounced transcription is possible in the absence of moderate H3K4me3. Shown are heat maps demonstrating high (blue) or low (yellow) levels around the TSS of both gene groups. Genes were ranked in both groups according to the level of H3K4me3 immediately downstream of the TSS.

**Figure S3.**
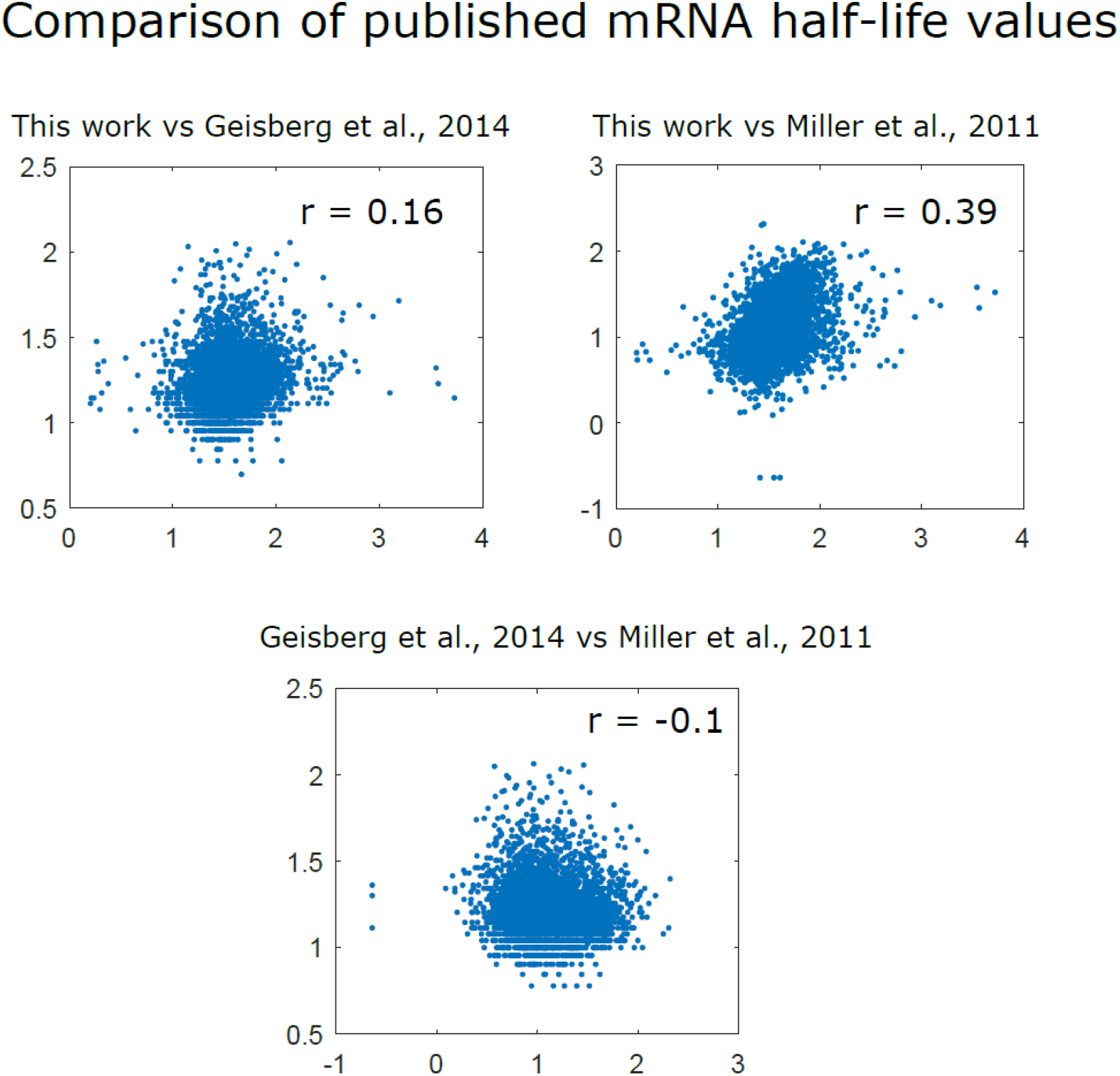
mRNA half-lives from this study and others correlate weakly to moderately with one another. Shown are scatter plots comparing the mRNA half-lives measured in this study and those indicated. r values shown are the Spearman correlation coefficients.

**Figure S4.**
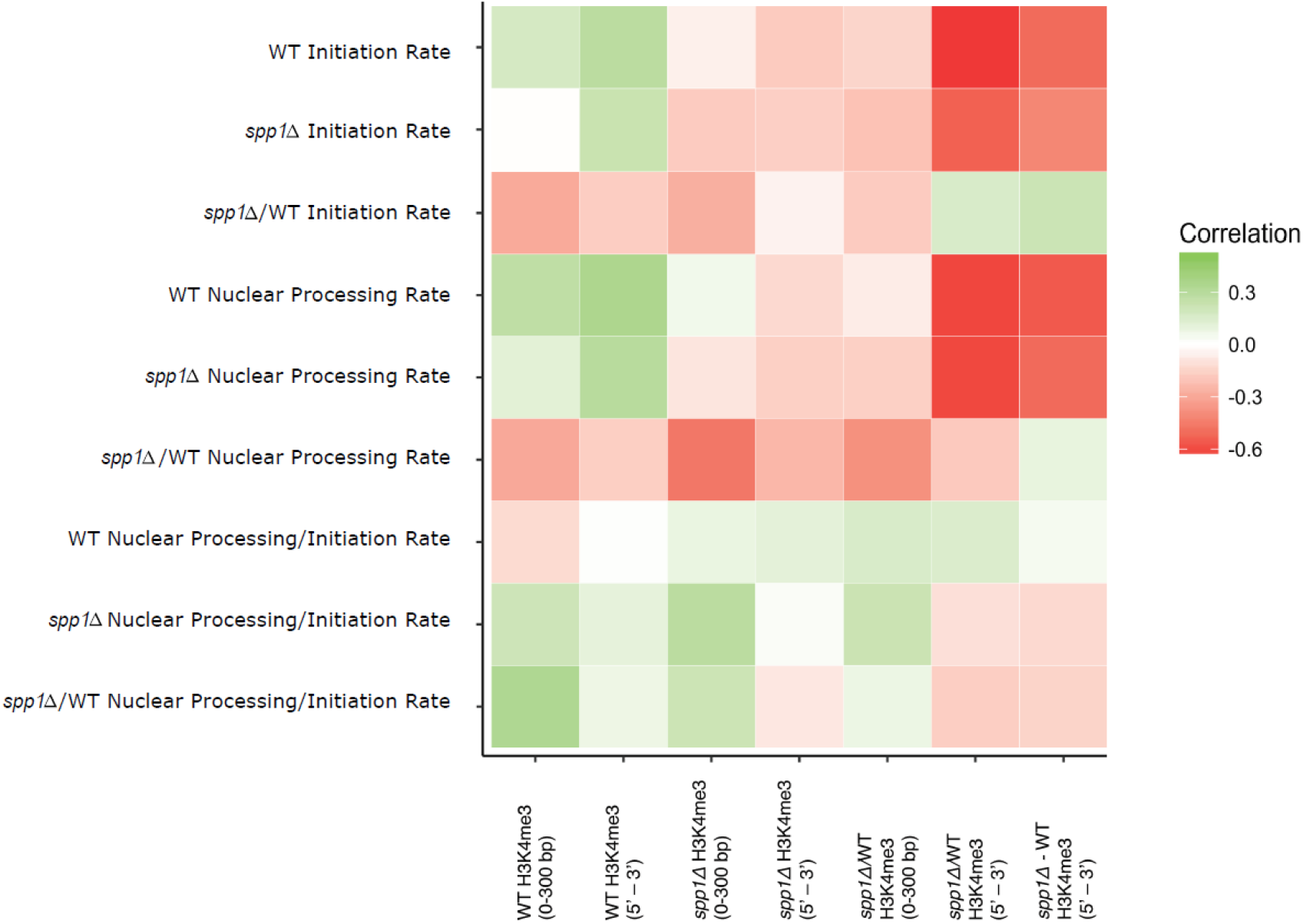
Correlation of transcription parameters derived from modelling with levels of H3K4me3. Shown are correlation coefficients obtained by comparing the estimated transcription parameters for the 12 genes studied by RNA-FISH with the levels of H3K4me3 in both wild-type and *spp1Δ* strains. H3K4me3 was averaged across either a 300 bp window immediately downstream of the TSS (0 – 300 bp) or the entire gene from TSS to poly(A) site (5’ to 3’).

## Notes

https://www.ncbi.nlm.nih.gov/geo/query/acc.cgi?acc=GSE133838

